## Supporting Information for "Sequence dependence of transient Hoogsteen base-pairing in DNA"

August 4, 2021

### 1 Adjustments to the general simulation protocol

For the outside path of sequence GAA, we average the metadynamics free-energy estimation from 1 to 6 ns. For the outside path of sequence CAG, we average the metadynamics free-energy estimation from 4 to 9 ns. For the outside path of sequence CAC, we average the metadynamics free-energy estimation from 1 to 4 ns. For both paths of sequence TAA and for the inside path of sequence CAT, we decrease the half-life to 10 ps, the attractor restraint force constant to 50 kcal/mol and the tube potential force constant to 20 kcal/mol. These changes are done either because the simulations crashed before the default time window (2 to 7 ns), or because they did not optimize to the desired pathway within the default time window with the general half-life and restraint parameters.

### 2 Collective variables

The collective variables (CVs) used in this work to bias the Watson-Crick-Franklin (WCF) to Hoogsteen (HG) base-pairing transition of the A16·T pair are:

- $\chi'$: The A16 base rolling torsion around the glycosidic bond defined by the pseudo-dihedral angle formed by the axis C1'-N9 and the vectors P2-P1 and P5-P6. Taken from [2] and shown in Figure S3A.
- $\theta$: The A16 base flipping torsion defined by the pseudo-dihedral angle (P1+P2)-P3-P4-P5. Taken from [3] and shown in Figure S3A.

Additionally, we analyze the following CVs describing different structural features of DNA, taken from [2]:

- $d_{\text{WCF}}$: The distance of the characteristic WCF H-bond between A16 (N1) and T9 (N3), shown in Figure S3B.
- $d_{\text{HG}}$: The distance of the characteristic HG H-bond between A16 (N7) and T9 (N3), shown in Figure S3B.
- $d_{\text{HB}}$: The distance of the conserved H-bond, present both in WCF and in HG, between A16 (N6) and T9 (O4), shown in Figure S3B.
- $d_{\text{CC}}$: The distance between A16 (C1') and T9 (C1'), shown in Figure S3B.
- $d_{\text{NB}}$: The distance between the neighboring bases of A16, described by the centers of mass P1' and P2', shown in Figure S3A.

*

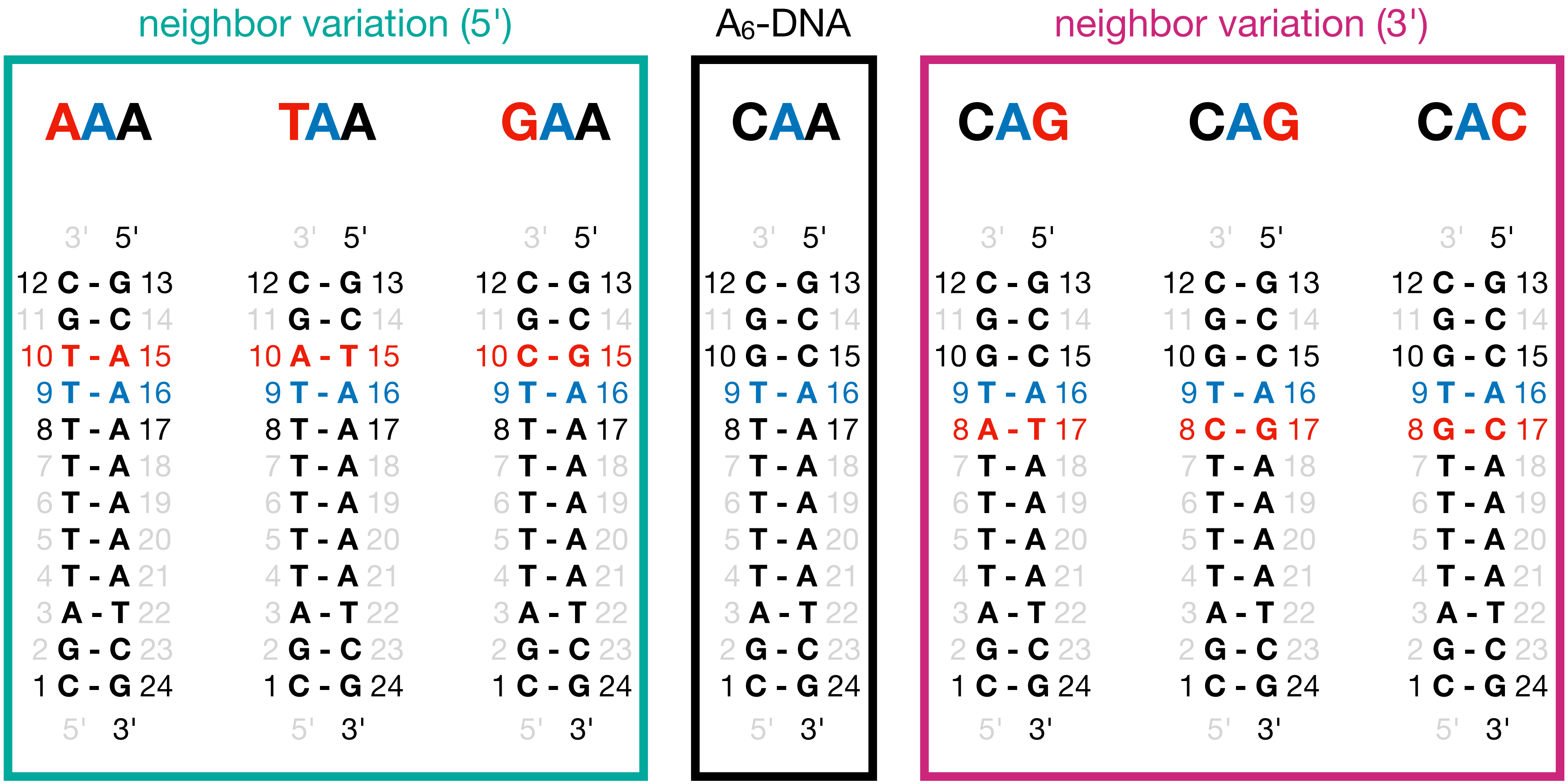


Figure S1: DNA sequence variations for which the transition from WCF to HG base pairing of A16·T9 is studied. The transitioning base pair A16·T9 is shown in blue. The original sequence $A_6$-DNA from [1] is outlined in black, and has the local environment CAA. Variations of the direct neighbor of A16 in the 5’ direction are outlined in teal, and have the local environments AAA, TAA and GAA. Variations of the direct neighbor of A16 in the 3’ direction are outlined in magenta, and have the local environments CAT, CAG and CAC.

Table S1: Stable-state characterization based on averages and standard deviations of several CVs during equilibrations at the WCF and the HG states for the sequences depicted in Fig. S1. The CVs—defined in [2]—describe the following structural changes. $\chi'$: A16 rolling angle; $\theta$: A16 flipping angle; $d_{\mathrm{WCF}}$: distance of the characteristic WCF hydrogen-bond; $d_{\mathrm{HG}}$: distance of the characteristic HG hydrogen-bond; $d_{\mathrm{HB}}$: distance of the conserved hydrogen-bond in both states; $d_{\mathrm{CC}}$: distance between the backbone C1' atoms of A16 and T9; $d_{\mathrm{NB}}$: distance between the neighboring nucleotides of A16.

| CV | state | AAA | TAA | GAA | CAA | CAT | CAG | CAC |
|---|---|---|---|---|---|---|---|---|
| $\chi'$ | WCF | 1.6±0.17 | 1.5±0.18 | 1.6±0.17 | 1.5±0.15 | 1.5±0.15 | 1.4±0.15 | 1.4±0.15 |
| (rad) | HG | -1.5±0.19 | -1.8±0.19 | -1.6±0.18 | -1.7±0.17 | -1.7±0.17 | -1.7±0.17 | -1.8±0.17 |
| $\theta$ | WCF | -0.1±0.09 | -0.1±0.12 | -0.1±0.08 | -0.1±0.11 | -0.1±0.09 | -0.1±0.11 | -0.1±0.09 |
| (rad) | HG | 0.0±0.05 | -0.0±0.08 | -0.0±0.06 | -0.0±0.05 | -0.0±0.05 | -0.0±0.06 | -0.0±0.06 |
| $d_{\mathrm{WCF}}$ | WCF | 3.0±0.12 | 3.0±0.12 | 3.0±0.13 | 3.0±0.12 | 3.0±0.12 | 3.0±0.13 | 3.0±0.13 |
| (Å) | HG | 5.8±0.21 | 6.0±0.58 | 6.0±0.48 | 5.9±0.19 | 6.0±0.24 | 6.0±0.28 | 6.0±0.32 |
| $d_{\mathrm{HG}}$ | WCF | 6.4±0.15 | 6.4±0.14 | 6.4±0.15 | 6.4±0.14 | 6.4±0.14 | 6.4±0.15 | 6.5±0.15 |
| (Å) | HG | 3.1±0.2 | 3.3±0.66 | 3.2±0.55 | 3.2±0.49 | 3.1±0.35 | 3.1±0.43 | 3.2±0.57 |
| $d_{\mathrm{HB}}$ | WCF | 3.0±0.18 | 3.0±0.2 | 3.0±0.19 | 3.0±0.17 | 3.0±0.2 | 3.0±0.2 | 2.9±0.16 |
| (Å) | HG | 2.9±0.22 | 3.0±0.71 | 3.0±0.52 | 3.0±0.42 | 2.9±0.16 | 2.9±0.18 | 2.9±0.23 |
| $d_{\mathrm{CC}}$ | WCF | 10.6±0.28 | 10.6±0.29 | 10.7±0.29 | 10.6±0.29 | 10.6±0.28 | 10.7±0.31 | 10.8±0.3 |
| (Å) | HG | 9.0±0.33 | 9.2±0.6 | 9.2±0.64 | 9.0±0.35 | 9.1±0.56 | 9.1±0.69 | 9.3±0.83 |
| $d_{\mathrm{NB}}$ | WCF | 7.5±0.31 | 7.9±0.4 | 7.4±0.3 | 7.9±0.36 | 7.6±0.44 | 7.8±0.41 | 7.8±0.41 |
| (Å) | HG | 7.6±0.31 | 7.9±0.4 | 7.4±0.29 | 7.8±0.38 | 7.7±0.36 | 7.9±0.4 | 8.0±0.4 |

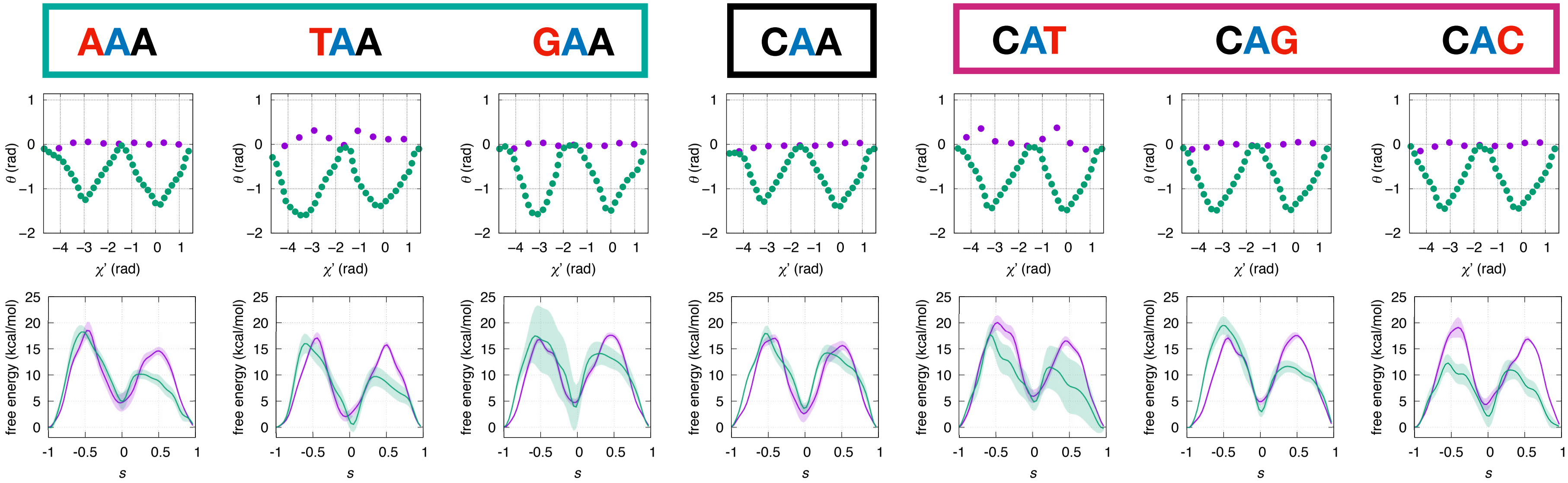


Figure S2: Transition pathways (top) and free-energy profiles (bottom) for the inside and outside mechanisms of WCF to HG base-pairing with local sequence variations. Sequences are as shown in Fig. S1. Variations of the direct neighbor of A16 in the 5' direction are outlined in teal, while in the 3' direction they are outlined in magenta. Inside paths are depicted in purple and outside paths in green. Sampling follows the scheme presented in Figure 1E in the main text.

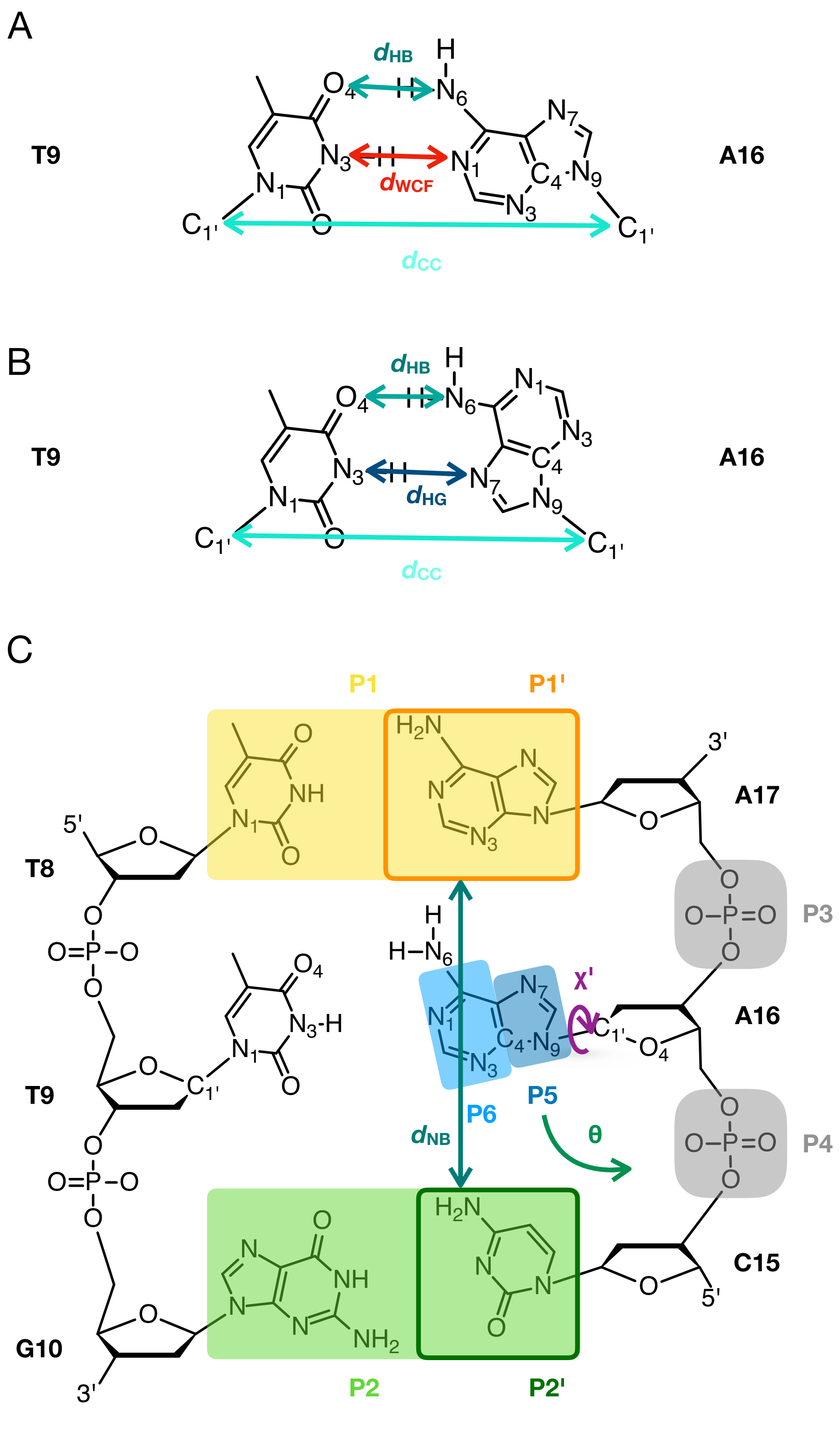


Figure S3: Scheme of the collective variables (CVs) biased or analyzed during the Watson-Crick-Franklin (WCF) to Hoogsteen (HG) base pairing transition. **A**: WCF base pair with graphical representations of the CVs: $d_{\mathrm{WCF}}$, $d_{\mathrm{HB}}$ and $d_{\mathrm{CC}}$. and HG base pair with graphical representations of the CVs: $d_{\mathrm{HG}}$, $d_{\mathrm{HB}}$ and $d_{\mathrm{CC}}$ (bottom); **B**: HG base pair with graphical representations of the CVs: $d_{\mathrm{HG}}$, $d_{\mathrm{HB}}$ and $d_{\mathrm{CC}}$ (bottom). **C**: A16·T9 bp and its two neighboring bps with graphical representations of the centers of mass involved in the calculation of CVs: $\chi'$, $\theta$ and $d_{\mathrm{NB}}$;
